# Imaging flow cytometry challenges the usefulness of classically used EV labelling dyes and qualifies that of a novel dye, named Exoria™ for the labelling of MSC-EV preparations

**DOI:** 10.1101/2021.06.09.447567

**Authors:** Tobias Tertel, Melanie Schoppet, Oumaima Stambouli, Ali Al-Jipouri, Patrick F. James, Bernd Giebel

## Abstract

Extracellular vesicles (EVs) are involved in mediating intercellular communication processes. An important goal within the EV field is the study of the biodistribution of EVs and the identification of their target cells. Considering that EV uptake is central for mediating the EVs role in intercellular communication processes, labelling with fluorescent dyes has emerged as a broadly distributed strategy for the identification of the EVs target cells and tissues. However, the accuracy and specificity of commonly utilized labelling dyes has not been sufficiently analyzed. By combining recent advancements in imaging flow cytometry for the phenotypic analysis of single EVs and aiming to identify target cells for EVs within therapeutically relevant MSC-EV preparations, we explored the EV labelling efficacy of various fluorescent dyes, specifically of CFDA-SE, Calcein AM, PKH67, BODIPY-TR-Ceramide and a novel lipid dye named Exoria. Our analyses qualified Exoria as the only dye which specifically labels EVs within our MSC-EV preparations. Furthermore, we demonstrate Exoria labelling does not interfere with the immunomodulatory properties of the MSC-EV preparations as tested in a multi-donor mixed lymphocyte reaction assay. Within this assay, labelled EVs were differentially taken-up by different immune cell types. Overall, our results qualify Exoria as an appropriate dye for the labelling of EVs derived from our MSC-EV preparations, this study also demonstrates the need for the development of next generation EV characterization tools which are able to localize and confirm specificity of EV labelling.

## Introduction

Extracellular vesicles (EVs) are membrane-enclosed particles in the nano- and micrometer range that are secreted into their extracellular environment by virtually all cells. According to their origin EVs are classified into different groups. The most prominent groups are exosomes, derivatives of the endosomal system with size ranges of 70-150 nm, microvesicles, shed offs of the plasma membrane of 100-1,000 nm, and apoptotic vesicles that can be as small as exosomes and as apoptotic bodies can reach sizes up to several micrometers (Raposo and Stoorvogel, 2013).

Despite these classes, EVs of each given subtype are very heterogeneous as well. Depending on the cell source they are originating from, they provide specific molecular compositions, qualifying them as a new class of biomarkers. Specifically, EVs residing in the plasma are increasingly used as biomarkers for different diseases (Fais et al., 2016; König et al., 2018; Vacchi et al., 2020). In addition to conventional methods, such as cytokine assays or cellular analyses, the prevalence of selected EV subpopulations can provide important new information on the course of respective diseases.

It became evident that EVs are of physiological relevance and mediate complex intercellular interactions at local and remote sites, both under healthy and pathological conditions (Yanez-Mo et al., 2015). Thus, it is a goal of many EV researchers to dissect such intercellular communication processes in a magnitude of different biological processes. In this context it is a relevant task to identify EV target cells. Addressing this challenge, it evolved as a common strategy to use fluorescent dyes considered to specifically label EVs and to apply labelled EV fractions either to assumed target cells/tissues *in vitro* or apply them *in vivo*. Among the commonly used dyes are dyes immediately integrating into membranes such as PKH dyes, dyes which become fluorescent after enzymatic reactions like non-fluorescent carboxyfluorescein diacetate succinimidyl ester (CFDA-SE) which by esterase is processed into carboxyfluorescein succinimidyl ester (CFSE), Calcein acetoxymethyl (Calcein AM) which binds calcium cations to become fluorescent or 4,4-difluoro-4-bora-3a,4a-diaza-s-indacene (BODIPY) conjugated fatty acids, e.g. ceramide (Chuo et al., 2018; Gray et al., 2015; Laulagnier et al., 2005; Nazarenko et al., 2013; Pospichalova et al., 2015; Pužar Dominkuš et al., 2018). Due to their micelle forming capabilities, use of several of these dyes are challenging for the EV field. Sophisticated preparation and washing procedures need to be followed to efficiently deplete dye aggregates, very often resulting in minor recovery rates (Dehghani et al., 2020). Furthermore, some dyes such as CFDA-SE require specific enzymatic activities to become fluorescent, in this case an esterase, to bind EV associated proteins and thus to efficiently label EVs (Banks et al., 2013).

For the quality control of dye labelled EV fractions, particle quantification methods are commonly used, and often performed by nanoparticle tracking analysis (NTA) or resistive pulse sensing. In 2011, our group in addition to Dragovic and co-workers introduced NTA as an “exosome” quantification method (Dragovic et al., 2011; Sokolova et al., 2011). However, the detailed comparison of data recorded using NTA and imaging flow cytometry (IFCM) (a more advanced EV characterization method), indicated that particle quantification methods are not appropriate to calculate EV concentrations in EV samples - unless they are ultra-pure (Droste et al., 2021). This cannot be achieved with conventional EV preparation techniques such as differential centrifugation, polymer precipitation or simple size exclusion technologies (Droste et al., 2021; Karimi et al., 2018; Vergauwen et al., 2017). In their traditional form, particle quantification methods cannot distinguish prepared particles such as protein precipitates, salt crystals and lipoprotein agglomerates from EVs. In addition, light scattering based methods can only detect some of the particles smaller than 100 nm due to their limited sensitivity (Giebel and Helmbrecht, 2017; van der Pol et al., 2014). Provided EVs are fluorescently labelled, IFCM grants an advanced platform for single EV detection (Görgens et al., 2019). Recently, we have optimized antibody labelling protocols for single EV analysis (Görgens et al., 2019; Tertel et al., 2020a; Tertel et al., 2020b). These protocols allowed us to investigate whether tetraspanins, specifically CD9, CD63 and CD81, whose expression within EV samples has been confirmed by WB, are co-localized on individual EVs or are recovered on distinct EV subsets. With this technology we analyzed EV preparations from mesenchymal stromal cell (MSC) conditioned, human platelet lysate supplemented media, whose therapeutic activities we study in different animal models and confirmed their therapeutic potential in a GvHD patient (Doeppner et al., 2015; Drommelschmidt et al., 2017; Kaminski et al., 2020; Kordelas et al., 2014; Ophelders et al., 2016; Wang et al., 2020). We demonstrated that CD9 and CD81 reside on different EV subpopulations all in the exosomal size range (Görgens et al., 2019). Notably, these results had been confirmed by an advanced multiplex bead-capturing procedure (Wiklander et al., 2018). Considering the method of IFCM as very informative for the EV characterization and intending to qualify a pan-EV labelling dye, we thus decided to evaluate the EV labelling efficacy of different dyes being used for EV marking. In addition to conventionally used BODIPY-TR-CER, Calcein AM, CFSE and PKH67 dyes, we included a novel dye named Exoria in our studies. Exoria, developed at Exopharm Ltd, was designed to be a pH stable fluorescence dye with reduced micelle forming propensity, which could incorporate into EVs.

In this study, the labelling efficiency of MSC-EV preparations with the dyes listed above was investigated. Counterstaining of PKH67 and Exoria labelled objects was performed with anti-tetraspanin antibodies. The impacts of Exoria labelling on the immunomodulatory capabilities of the MSC-EV preparation were investigated in a multi-donor mixed lymphocyte reaction (mdMLR) assay. Furthermore, the uptake of Exoria labelled objects by the different immune cells within the mdMLR assay were documented.

## Material and methods

### Preparation of EVs from MSC conditioned cell culture media

MSC-EVs were prepared form human platelet lysate containing MSC-conditioned media by polyethylene glycol 6000 precipitation followed by ultracentrifugation, as described previously (Borger et al., 2020; Kordelas et al., 2014; Ludwig et al., 2018). Conditioned media were harvested every 48 h. Obtained MSC-EV preparations were diluted in NaCl-HEPES buffer (Sigma-Aldrich, Taufkirchen, Germany) such that 1 mL of final samples contained the preparation yield of the conditioned media of approximately 4.0 × 10^7^ cells.

### Characterization of the EV preparations

Obtained EV preparations were characterized according to the MISEV criteria (Thery et al., 2018). Briefly, average particle concentrations were determined by NTA on a ZetaView PMX-120 platform equipped with the software version 8.03.08.02 (ParticleMetrix, Meerbusch, Germany) as described previously (Ludwig et al., 2018). Protein concentration was determined by bicinchoninic acid (BCA) assay (Pierce, Rockford, IL, USA) in 96-well plates according to the manufacturer’s recommendations. The presence of EV specific proteins (CD9, CD63, CD81 and Syntenin) and the absence of impurities (Calnexin) were confirmed in Western Blots performed as described previously (Ludwig et al., 2018).

### EV labelling with different dyes

The staining with CFSE (Thermo Fisher Scientific, Darmstadt, Germany) was based on the manufacturer’s protocol. Slight modifications were required to reduce the signal from unbound CFSE. Briefly, the CFSE stock solution was diluted to a working solution of 10 µM CFSE. The solution was centrifuged three times for 10 min at 17,000 *x g* (see also supplement figure 1). Subsequently, 25 µL of MSC-EV preparations, corresponding to the amount of EVs derived from 1×10^6^ MSCs, were incubated with the centrifuged CFSE solution for 20 min at 37°C. The sample was diluted 1:20 to a final volume of 1 mL prior to analysis.

The staining of MSC-EV preparations with Calcein AM followed the manufacturer’s instructions. Briefly, 25 µL of MSC-EV preparations were incubated with 25 µL of a 20 µM solution of Calcein AM (Thermo Fisher Scientific) for 40 min at 37°C. The sample was diluted 1:20 to a final volume of 1 mL to reduce background noise, avoiding the requirement of a washing step.

The staining of MSC-EV preparations with BODIPY-TR-Ceramide followed the manufacturer’s instructions. Briefly, 25 µL of the MSC-EV preparation, corresponding to EVs purified from 4×10^6^ MSCs, were incubated with 25 µL of a 20 µM solution of BODIPY TR Ceramide (Thermo Fisher Scientific) for 20 min at 37°C. 450 µL of 0.9% NaCl with 10mM HEPES (0.9% NaCl, Melsungen, B. Braun; HEPES, Thermo Fisher Scientific) buffer was added and the EVs were washed by using a Centrifugal Concentrator (Vivaspin 500; Sartorius, Göttingen, Germany). The retentate was adjusted to 500 µL prior to analysis.

The staining of MSC-EV preparations with PKH67 followed the manufacturer’s instructions for labeling EVs (Thermo Fisher Scientific). Briefly, using 200 µL of given MSC-EV preparations, corresponding to EVs purified from 8×10^6^ cells, the solution was adjusted with Diluent C to a final volume of 1 mL. 6 µL of PKH67 dye was added to each tube and mixed continuously for 30 seconds. After 5 min at room temperature, the solution was quenched by adding 2 mL of 10% (w/v) bovine serum albumin fraction 5 (Carl Roth, Karlsruhe, Germany). Serum-free medium, DMEM low glucose (PAN Biotech, Aidenbach, Germany) supplemented with 100 U/ mL penicillin-streptomycin-glutamine (Thermo Fisher Scientific, Darmstadt, Germany), was used to adjust the volume to 8.5 mL. 1.5 mL of a 0.971 M sucrose solution (Carl Roth) was added to the bottom of the tube, and the tube was centrifuged for 2 hours at 190,000 *x g* in a swing-out rotor (SW40 Ti; Beckman Coulter, Krefeld, Germany; k-factor: 137) at 4 °C. The supernatant was discarded, and the pellet resuspended in Na-HEPES buffer. After resuspension, the volume was adjusted to 5 mL and transferred to a Centrifugal Concentrator (Vivaspin 6; Sartorius). The retentate was adjusted to 120 µL prior to analysis.

The MSC-EV preparations were stained with Exoria following the protocol provided by Exopharm Ltd. Exoria was provided as a lyophilized powder. 1 mg was resuspended with 1 mL buffer to a final concentration of 0.2 µM. Like CFDA-SE, the Exoria solution was centrifuged for 10 min at 17,000 *x g* to reduce background noise to a minimum. Briefly, for the EV-labelling 25 µL of the MSC-EV preparations were incubated with 25 µL of a prepared, centrifuged Exoria solution (0.2 µM) for 1 hour at 37°C. The sample was diluted 1:20 to a final volume of 1 mL prior to analysis. For EV uptake experiments Exoria labelled MSC-EV preparations were cleared from EV unbound Exoria by ultrafiltration. Briefly, after labelling with Exoria, the MSC-EVs were washed by via centrifugation at 12,000xg through Vivaspin 500 filters (Sartorius) for 10 min. The retentate was collected as labelled EV sample.

### Antibody labelling of prepared EVs

After dye labelling, 5 µL of Exoria stained MSC-EV samples were mixed with 20 µL of a 10 nM anti-human CD9 FITC (EXBIO, Vestec, Czech Republic), 12 nM anti-human CD63 AF488 (EXBIO) or 13 nM anti-human CD81 FITC (Beckman Coulter) antibody solution, respectively. For PKH67 stained MSC-EV samples, incubated with 10 nM anti-human CD9 PE (EXBIO), 12 nM anti-human CD63 PE (EXBIO) or 13 nM anti-human CD81 PE (Beckman Coulter), respectively, for 2 hours at room temperature as described previously (Tertel et al., 2020b). Accordingly, isotype controls were performed (see also supplement table 1) For Exoria, final preparations were diluted to 500 µL for CD9 (end dilution factor of 1 to 100) and 200 µL for CD63 and CD81 analyses (end dilution factor of 1 to 40). The preparations for PKH67 were diluted to 100 µL for all three analyses (1:20 dilution).

### Detergent control

To test for the EV nature of labelled objects detergent controls were performed by adding a sample volume of a 2% (w/v) NP-40 solution (Calbiochem, San Diego, CA, USA) to the samples.

### IFCM analyses

All samples were measured using the built-in autosampler from U-bottom 96-well plates (Corning Falcon, cat 353077) with 5 min acquisition time per well on the AMNIS ImageStreamX Mark II Flow Cytometer (AMNIS/Luminex, Seattle, WA, USA). All data were acquired at 60x magnification at low flow rate (0.3795 ± 0.0003 μL/min) and with removed beads option deactivated as described previously (Görgens et al., 2019; Tertel et al., 2020a). The data was analyzed as described previously (Tertel et al., 2020b). Additional settings can be found in supplement table 2 and 3.

### Multi-donor mixed lymphocyte reaction (mdMLR)

The immunomodulatory potential of Exoria labelled and non-labelled MSC-EV preparations were compared in a multi-donor mixed lymphocyte reaction assay (MLR) exactly as described previously (Madel et al., 2020). Briefly, Ficoll prepared peripheral blood mononuclear cells (PBMC) of 12 healthy donors were mixed in equal proportions, aliquoted and stored in the vapour phase of liquid nitrogen until usage. After thawing 600,000 cells were seeded per well of a 96-well U-bottom shape plates (Corning, Kaiserslautern, Germany) and cultured in 10% human AB serum (produced in house) and 100 U/mL penicillin and 100 µg/mL streptomycin (Thermo Fisher Scientific) supplemented RPMI 1640 medium (Thermo Fisher Scientific) in a final volume of 200 µL per well, either in the presence or absence of MSC-EV preparations to be tested. After 5 days, cells were harvested, stained with a collection of different fluorescent labelled antibodies (CD4-BV785; BioLegend, San Diego, CA, USA; CD25-PE-Cy5.5; BD Bioscience; and CD54-AF700; EXBIO) and analysed on a Cytoflex flow cytometer (Software CytExpert 2.3, Beckman-Coulter). Activated and non-activated CD4^+^ T cells were discriminated by means of their CD25 and CD54 expression, respectively. Typically, 5 µL of MSC-EV preparations to be tested were applied into respective wells. The following antibodies were used to further discriminate subpopulations: CD8-BV650 (BioLegend), CD14-PO (EXBIO), CD19-ECD (Beckman Coulter) and CD56-APC (BioLegend). The evaluation of the data was carried out with the Kaluza software (Version 2.1, Beckman Coulter).

### Statistics

The statistics and graphical presentation were performed with GraphPad version 8.4.3. Mean values ± standard deviation are provided.

## Results

### CFSE, Calcein AM and BODIPY-TR-Ceramide do not label MSC-EVs

Aiming to identify a dye allowing specific labelling of EVs in therapeutically active MSC-EV preparations, we decided to evaluate the accuracy of conventionally used EV labelling dyes, specifically CFSE, Calcein AM, PKH67, BODIPY-TR-Ceramide and a novel lipid dye named Exoria. MSC-EV preparations that have been extensively explored in various animal models had been obtained from supernatants of MSCs raised in 10% human platelet lysate supplemented media by our well established PEG-ultracentrifugation protocol (Borger et al., 2020; Doeppner et al., 2015; Drommelschmidt et al., 2017; Gussenhoven et al., 2019; Kaminski et al., 2020; Kordelas et al., 2014; Ludwig et al., 2018; Ophelders et al., 2016; Wang et al., 2020). Since micelle formation of some of the dyes have been reported and following the MIFlowCyt-EV recommendation (Welsh et al., 2020), we initially added all of the labelling dyes but the EV sample to the NaCl-HEPES buffer, the buffer MSC-EVs are suspended in. Samples were processed according to the manufacturer’s recommendation and analyzed by IFCM with protocols that we have successfully established for the characterization of antibody labelled MSC-EVs (Görgens et al., 2019; Tertel et al., 2020a; Tertel et al., 2020b).

Notably, depending on the manufacturer’s protocol, different amounts of MSC-EV preparation was required. For the CFSE, Calcein-AM, BODIPY-TR Ceramide-(BODIPY) and Exoria labelling we started with volumes of 25 µL of MSC-EV preparation, for PKH67 with 200 µL. Initially, we analyzed all recorded objects. Based on our prior experience, sEVs appear as fluorescently labelled objects with minimal side scatter signals (SSC). Upon comparing the dye only solutions, CFSE, Calcein AM and Exoria were observed to contain no objects. In contrast, upon analyzing the BODIPY and PKH67 solutions, solid populations of labelled objects with minimal side scatter signals were identified (Figure 1). This data implies micelle or aggregate formation of BODIPY and PKH dyes.

**Figure 1:**
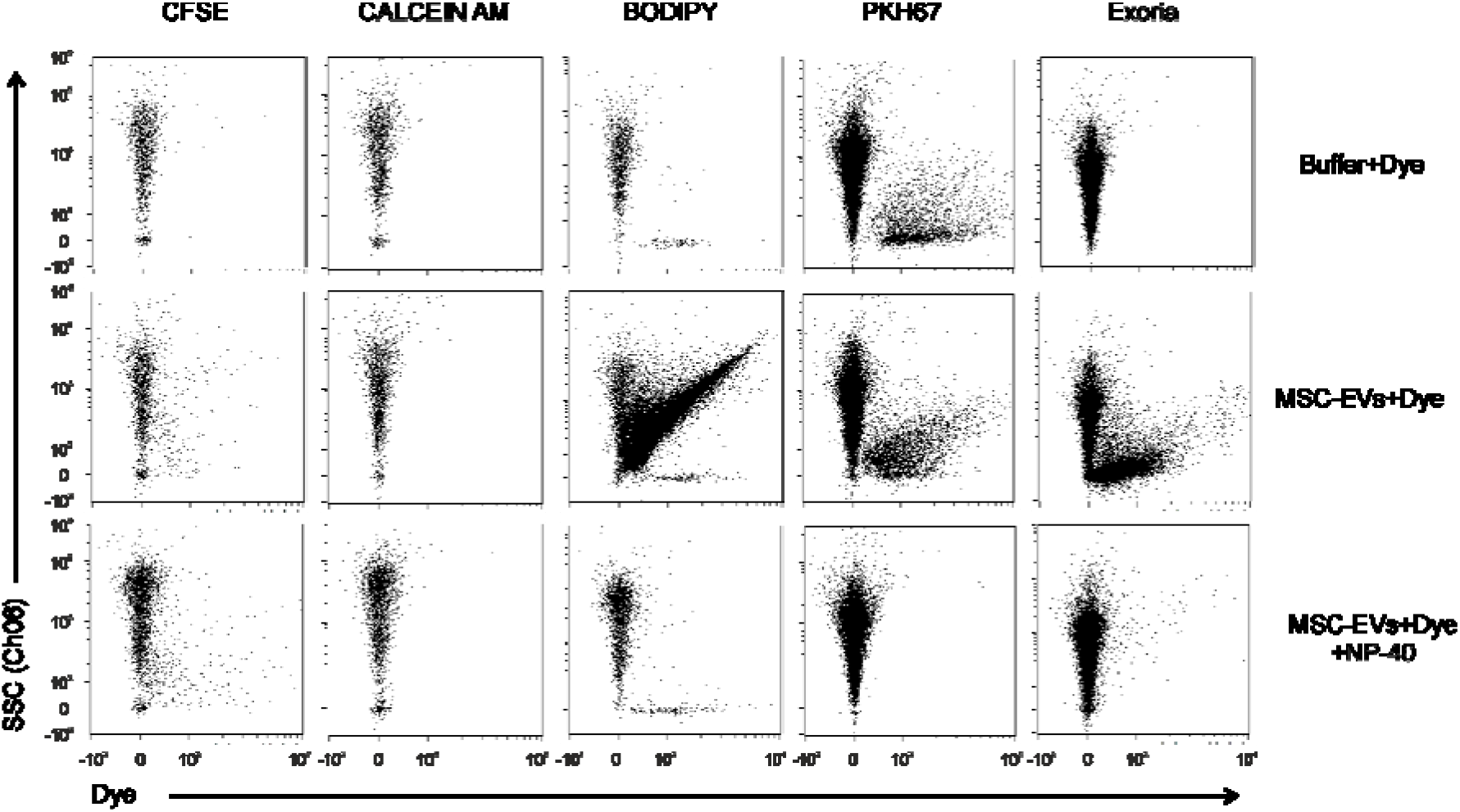
PKH67 and Exoria specifically label objects within MSC-EV preparation with sEV-like side scatter properties. Fluorescent labelling procedures were performed for CFSE, Calcein AM, BODIPY, PKH67 and Exoria in the absence of any EV preparation (upper row), and in the presence of MSC-EVs, alone (middle row) or in the presence of MSC-EVs and the detergent NP40. Fluorescence intensities of the dye labelled objects (x-axis) are plotted against the intensity of their size reflecting SSC signals (y-axis).

Subsequently, MSC-EV preparations labelled using the same procedure were analyzed. In contrast to the buffer only solutions, solid populations of labelled objects were observed after BODIPY, PKH67 and Exoria labelling and some objects following CFSE labelling (Figure 1). Calcein AM failed to label any detectable objects. BODIPY^+^ objects that were not recovered in the buffer-BODIPY solution control revealed side scatter signals that were much higher than those typically seen for small EVs (sEVs). In contrast, the light scattering properties of the objects specifically labelled with PKH67 or Exoria reflect those of sEVs. Notably, in good agreement with published reports that PKH dyes increased the size of labelled EVs (Dehghani et al., 2020; Morales-Kastresana et al., 2017a), the PKH67^+^ objects specifically labelled in the MSC-EV preparation indicated higher side scatter signals than Exoria^+^ objects (Figure 1).

To determine whether the specifically labelled objects are detergent-sensitive, the dye-labelled MSC-EV samples was treated with NP40. While the BODIPY^+^ objects specifically detected in MSC-EV preparations, those with the higher light scattering properties, and all PKH67^+^ and Exoria^+^ objects disappeared following NP40 treatment. In contrast, the population of CFSE^+^ objects and the BODIPY^+^ objects with sEV light scattering properties were hardly affected by the NP40 treatment. To this end, we considered neither detergent resistant CFSE^+^ nor the BODIPY^+^ objects with low light scattering properties as small EVs. Coupled to the failure of Calcein AM to label any specific objects, we excluded CFSE, Calcein AM and BODIPY from all later analyses and focused on exploring the accuracy of PKH67 and Exoria as MSC-EV labelling dyes.

### PKH67 fails to effectively label CD9^+^, CD63^+^ and CD81^+^ sEVs

Next, we investigated the potential co-localizations of PKH67 with known EV markers. To this end, we continued with the well characterized MSC-EV preparations. These were stained either by PKH67 alone or in combination with any of the following antibodies: anti-CD9, anti-CD81 or anti-CD63 antibodies. To reduce the background noise and to exclude coincident events, the simultaneous detection of two or more independent objects at the same time (coincidences with high object numbers), we applied an optimized gating strategy. Briefly, we focused on objects recognized as singlets in the PKH67 channel without a simultaneous antibody signal or as singlets in the antibody channel without a simultaneous PKH67 signal, and on events appearing in both channels as singlets not providing two individual objects (Figure 2A).

**Figure 2:**
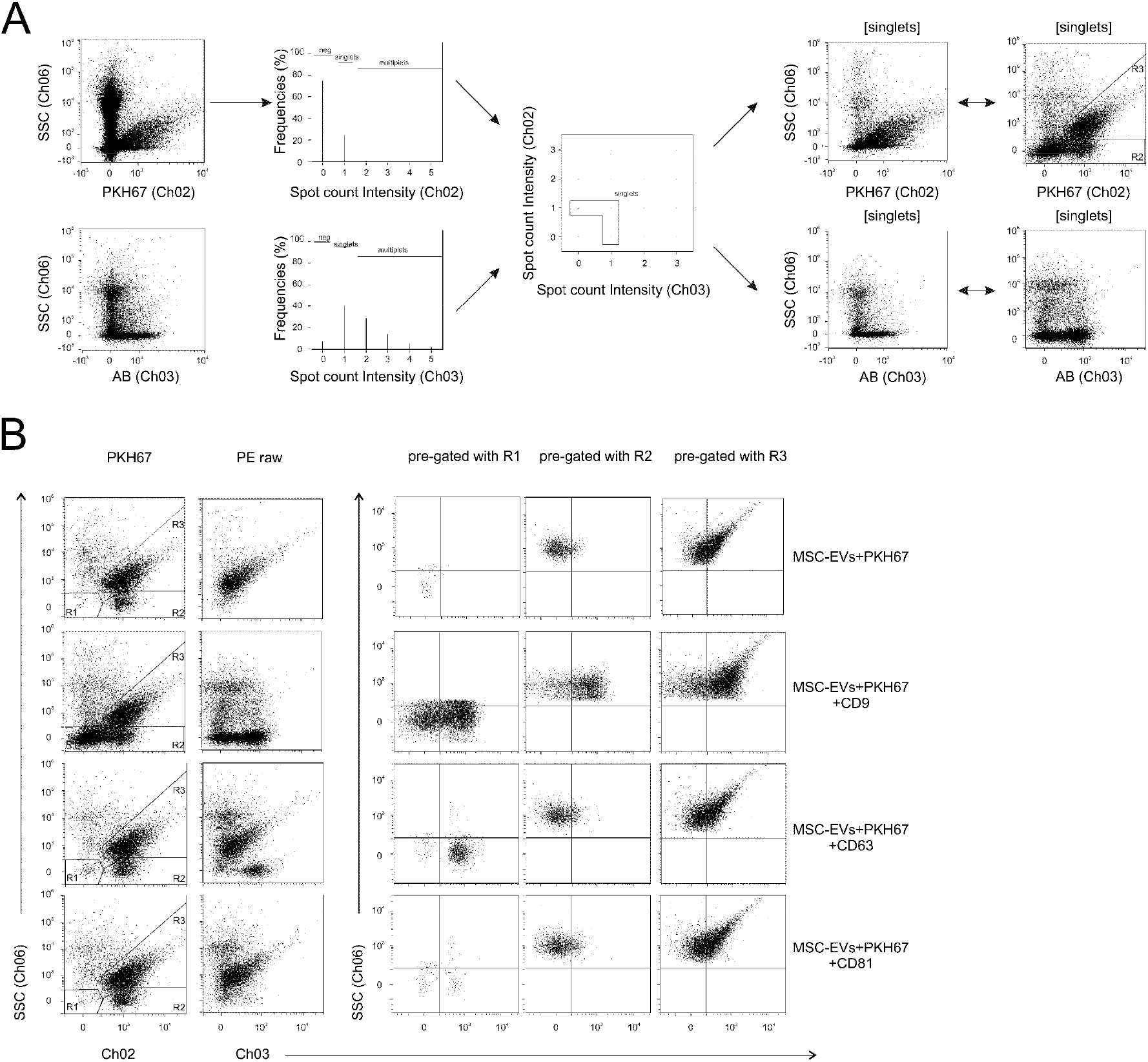
PKH67 fails to effectively label small MSC-EVs. (A) Gating strategy of the detected objects. Exemplarily, the gating strategy is presented for MSC-EV preparations counterstained with PKH67 and anti-CD9 antibodies. Both fluorescence channels (CH02 and CH03) are initially plotted against the side scatter intensities (SSC) of all recorded objects. Number of co-incident objects per channel are depicted (2^nd^ column). Of all recorded objects, the only objects considered in subsequent analyses were those that showed either single signals in the PKH67 or the antibody channel, or single signals in both channels (singlets). Within the CH02 SSC singlet plots three different gates were defined with singlets in R1 and R2 with low and in R3 with concrete side scatter signals. Objects in R1 revealed no PKH67 and those in R2 and R3 concrete PKH67 signals. Ch02 signals plotted against SSC signals of the singlets are shown as well, either in the same plot size as in the left column before gating or in the zoom in versions of the same plots (right column). (B) Distribution of recorded singlets in R1, R2 and R3 without antibody labelling or following anti-CD9, anti-CD63 or anti-CD81 labelling, respectively. Plotting of the fluorescence intensities of singlets in the PKH67 (Ch02) or the antibody channel (Ch03) against the singlets’ side scatter intensities. Column 3 to 5, fluorescence intensities of R1 to R3 gated singlets. (C) Number of events in gates R1-R3 for the respective measurements. The mean values ± standard deviation indicated.

Upon plotting side scatter against PKH67 intensities, many more objects were recovered in samples that had been counterstained by anti-CD9 antibodies than in the PKH67 labelled buffer and MSC-EV containing controls. Most of these objects were negative for PKH67 and showed low SSC signals. The region containing these objects was defined as R1. For anti-CD9 staining, 6672 ± 1170 objects were recovered in the region R1. Notably, hardly any objects were recovered in R1 in the PKH67 labelled MSC-EV samples that were not counterstained by antibodies. A slight increase in objects numbers was recorded when PKH67 labelled MSC-EV samples were counterstained with anti-CD63 (221 ± 42 objects) or anti-CD81 (96 ± 32 objects) antibodies. Most of the objects that were positive for PKH67 revealed solid SSC signals. The region including these objects was defined as R3. A smaller number of PKH67 labelled objects was identified with low SSC signals that was clustered in a region defined as R2. In contrast to the number of objects in R1, the numbers of objects in R2 and R3 were only slightly affected by the antibody labelling procedures (Figure 2B and 2C). Within the antibody non-labelled control 2040 ± 344 objects were recovered in R2, following anti-CD9 staining 2927 ± 466 objects, following anti-CD63 staining 1747 ± 141 objects and following anti-CD81 staining 1734 ± 200 objects. In all antibody labelled MSC-EV preparations more objects were found in R3 (anti-CD9: 6625 ± 803 objects; anti-CD63: 5915 ± 271 objects; anti-CD81: 5777 ± 675 objects) than in the antibody non-labelled control (2046 ± 157 objects). To analyze objects within the 3 different regions in more detail, their antibody-labelling intensities were plotted against PHK67 labelling intensities. The results clearly confirm that a huge proportion of the objects in R1 were effectively labelled by anti-CD9 antibodies. Although the R1 object populations were much smaller following anti-CD63 and anti-CD81 than after anti-CD9 antibody staining, a proportion of these objects was clearly recognized as CD63^+^ or CD81^+^, respectively (Figure 2B). In contrast, all objects in R2 or in R3 appeared as CD63^-^ and CD81^-^ objects, most of which can be labelled by anti-CD9 antibodies. Notably, the frequencies of CD9^+^, CD63^+^ and CD81^+^ objects recovered in R1 are congruent to our previous observations that MSC-EV preparations contain a dominating CD9^+^CD81^-^ and a minor CD9^-^CD81^+^ sEV population (Görgens et al., 2019). Overall, we consider that most antibody stained objects in R1 were sEVs not being labelled by PKH67 and that most of the sEVs can only be detected if they are succesfully stained with any of the 3 antibodies. Thus, our data question PKH67 as efficient MSC-sEV labelling dye.

### Exoria effectively labels CD9^+^, CD63^+^ and CD81^+^ EVs in MSC-EV preparations

Next, the reliability of the Exoria as an EV labelling dye was investigated in a comparable manner to PKH67. To this end, the MSC-EV preparations (n=3) were either stained with Exoria alone or in combination with anti-CD9, anti-CD63 or anti-CD81 antibodies, respectively. Without defining R1-3 sub gates, gating strategies were applied as depicted in Figure 2A. In contrast to the PKH67 labelling experiments, many more objects with lower side scatter signal intensities were labelled by Exoria, even in the absence of any of the three different antibodies. No clear increase in the numbers of detected objects was observed between MSC-EV samples that were solely labelled by Exoria or in addition by anti-CD9 antibodies (Figure 3). Thus, in contrast to the PKH67 labelling, Exoria labelling is sufficient to label most of the sEVs within our MSC-EV preparations. Interestingly, upon plotting the Exoria labelling intensities against that of the different antibodies, it appears that CD81^+^ objects are more intensively labelled with Exoria than CD63^+^ objects. Furthermore, the results imply that more than 90% of the CD9^+^ and CD81^+^ objects had been labelled with Exoria but only 60% of the CD63^+^ EVs. All labelled objects were confirmed to be detergent sensitive (Suppl. Figure 2). Even though we do not have an explanation for the weaker Exoria stainability of CD63^+^ compared to CD9^+^ and CD81^+^ objects, overall, the data demonstrate that Exoria successfully labels most of the sEVs in our MSC-EV preparations.

**Figure 3:**
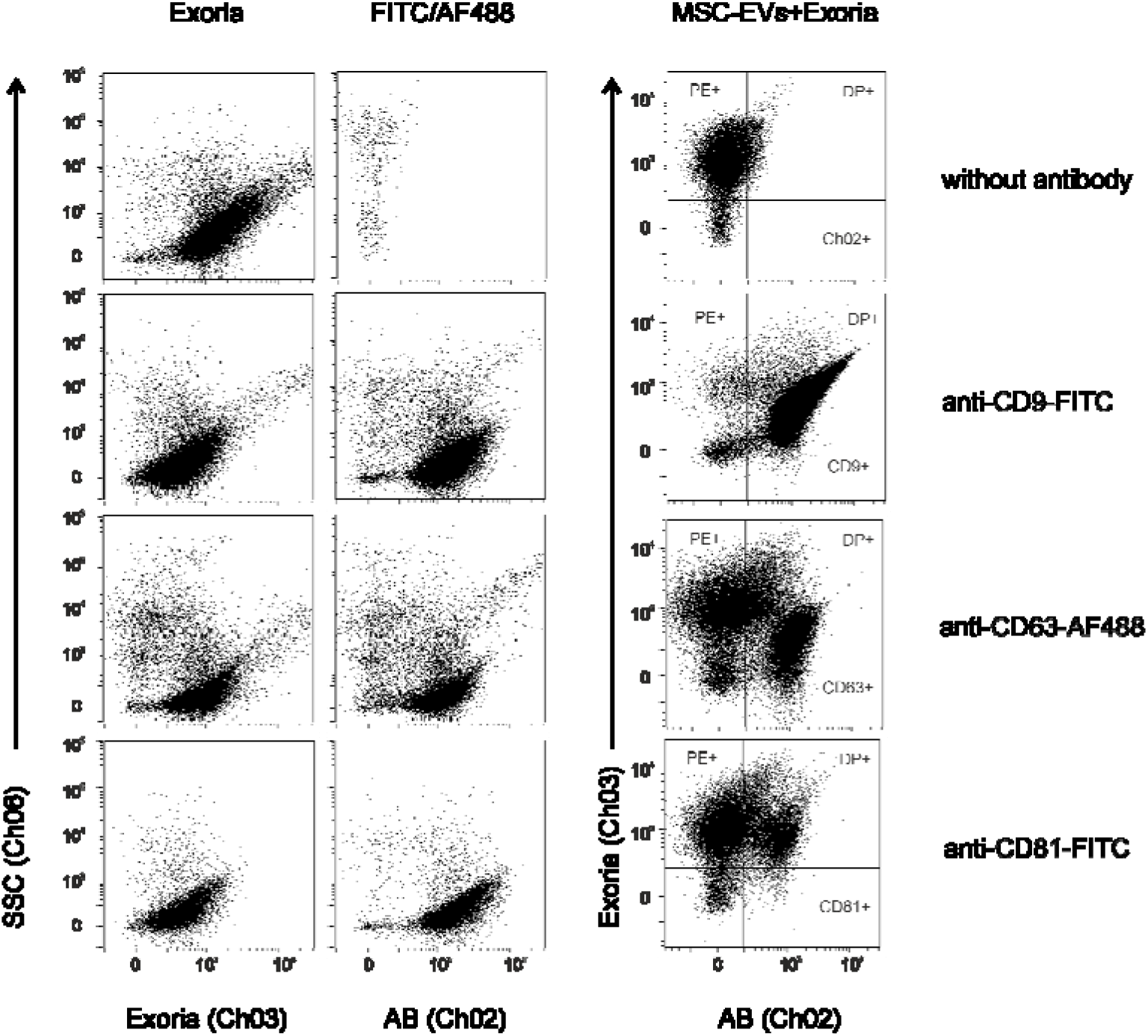
Exoria stains tetraspanin containing EVs. MSC-EV preparations were counterstained with Exoria and either anti-CD9, anti-CD63 or anti-CD81 antibodies. The same gating strategy as described in Figure 2 was applied. Fluorescence intensities of singlets are plotted against the side scatter (SSC) intensities either for the antibody (Ch02) or the Exoria (Ch03) channel. In the third column Exoria signals are plotted against the signals of respective antibodies. NP40 lysis controls are presented in Suppl. Figure 2.

### Exoria staining does not affect the immunomodulatory capacity of the MSC-EV preparations

Next, we investigated whether Exoria affects the MSC-EV preparation’s immunomodulatory capability. To this end, the activity of Exoria stained MSC-EV preparations were compared to corresponding, non-labelled MSC-EV preparations in a multi-donor mixed lymphocyte reaction (mdMLR) assay. Upon pooling of mononuclear cells from the peripheral blood of 12 different healthy donors (PBMCs), allogenic immune reactions are induced that can be monitored by the activation status of CD4^+^ T cells. Following 5 days in culture, approximately a quarter of all CD4^+^ T cells express the interleukin-2 receptor (CD25) and the intercellular adhesion molecule-1 (CD54), indicating T cell activation (Figure 4). As previously described, MSC-EV preparations with immunomodulatory capabilities effectively reduce the content of activated CD4^+^ T cells (Madel et al., 2020). Consistently, in the presence of the non-labelled MSC-EV preparations only 16% of the monitored CD4^+^ T cells were found to display the activation cell surface markers (Figure 4B). In the presence of Exoria labelled MSC-EV preparations (n=3) we observed a comparable reduction in CD4^+^ T cell activation (Figure 4B). Notably, Exoria itself did not influence the activation status of CD4^+^ T cells (Figure 4B). Thus, Exoria does not recognizably affect the immunomodulatory capability of the applied MSC-EV preparations.

**Figure 4:**
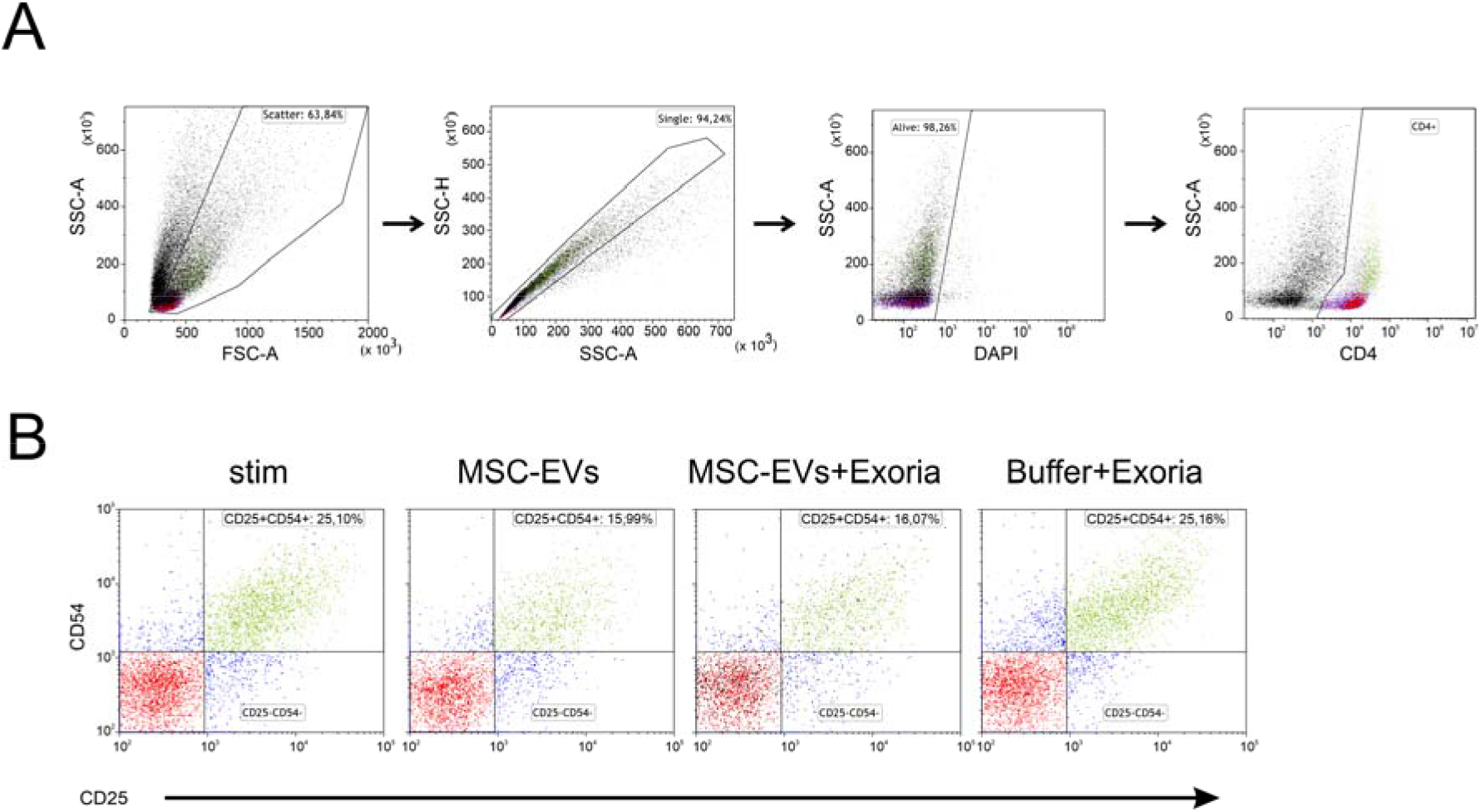
Exoria staining does not affect the immunomodulatory capability of MSC-EV preparations. Mixtures of PBMCs of 12 different donors were cultured in the presence or absence of non-labelled or Exoria labelled MSC-EV preparations, or in the presence of the Exoria dye for 5 days. Thereafter, cells were harvested and stained with DAPI and fluorescently labelled anti-CD4, anti-CD25 and anti-CD54 antibodies and analyzed by conventional flow cytometry. (A) Gating strategy for CD4 T cells. Living cells were identified according to their forward and side scatter features as singlets and DAPI negative cells. CD4 T cells were gated as CD4^+^ living cells. (B) Fluorescent intensities of CD25 and CD54 gated living CD4^+^ cells of mdMLR assays cultured in the absence of any additives (stim), in the presence of non-labelled MSC-EVs (MSC-EVs), Exoria labelled MSC-EVs (MSC-EVs +Exoria) or in the presence of buffer solved Exoria (Buffer + Exoria).

### Exoria stain MSC-EVs exhibit different uptake potential across immune cell subtypes of a mdMLR assay

To test whether Exoria EV labellinglabelling allows the identification of EV-up taking cells, we examined the labelled EV uptake of the different immune cells within the mdMLR assay, next. To this end, a pool of PBMCs derived from 12 healthy donors were cultured for 5 days in the presence of Exoria labelled MSC-EVs (n=3) that had been cleared from excessed Exoria dye by ultrafiltration. Thereafter, cells were harvested, antibody labelled and analyzed by flow cytometry. The content of Exoria labelled cells within different PBMC subtypes was determined. Almost all monocytes (CD14^+^ cells, 99%) revealed Exoria signals. In contrast, only proportions of the different lymphocytes appeared as Exoria positive cells, i.e. 71% of all CD4^+^ T cells (CD4^+^ cells), 34% of all CD8^+^ T cells (CD8^+^ cells), 72% of all B cells (CD19^+^ cells) and 15% of all NK cells (CD56^+^ cells). In addition, IFCM was used to visualize the subcellular staining of Exoria positive cells (Figure 5C). Obtained images reveal concrete labelled structures that according to our experience are located subcellularly. Thus, Exoria labelled EVs within our MSC-EV preparations are specifically taken up by different contents of the immune cell types within the assay.

**Figure 5:**
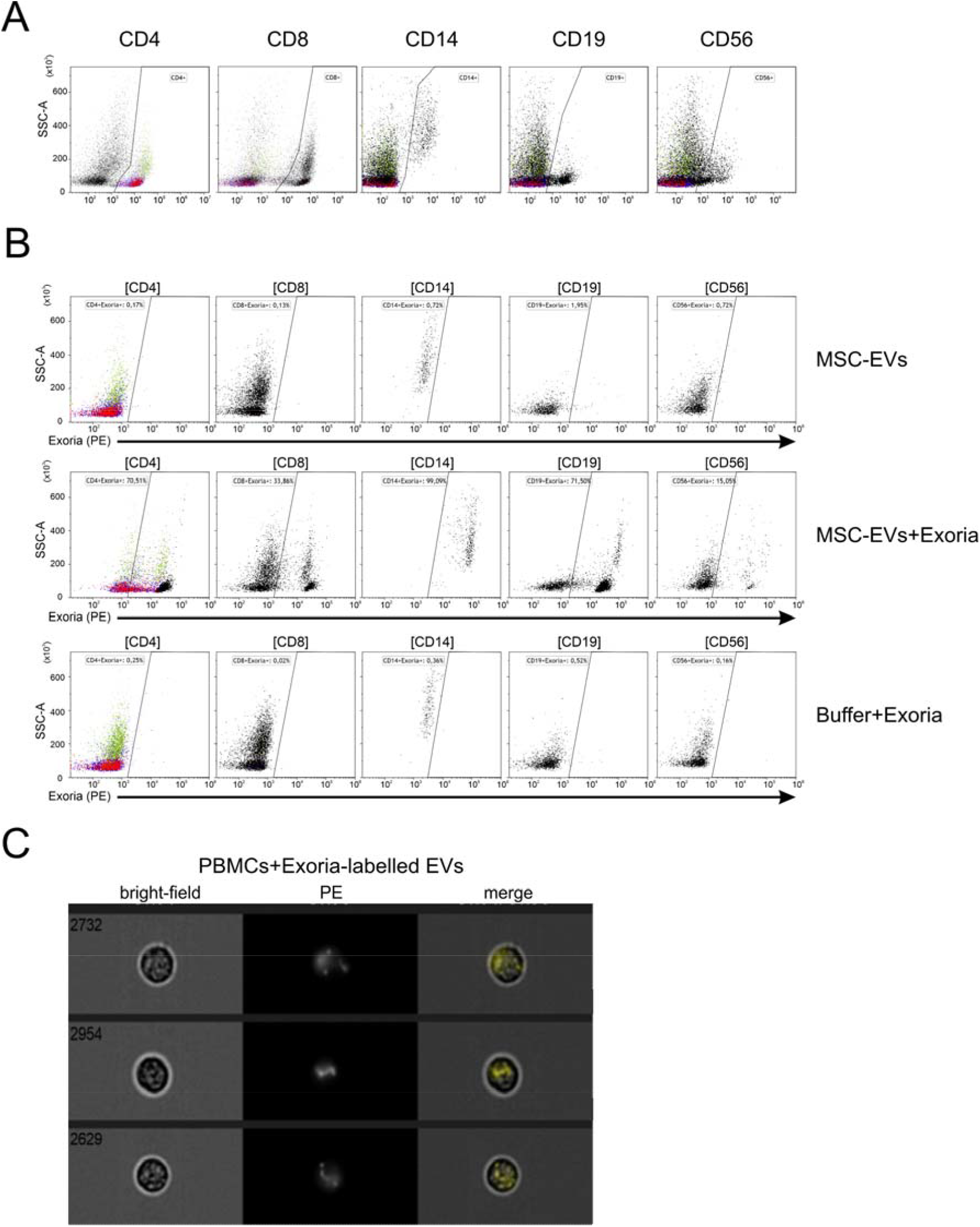
Analyses of the various immune cell types within the mdMLR assay reveal differences in uptake of Exoria-stained MSC-EVs. Immune cells of the mdMLR were examined for uptake of Exoria-stained MSC-EV preparations. (A) Discrimination of the different subpopulations was performed by using antibodies against CD4 and CD8 (T cells), CD19 (B cells), CD56 (NK cells), and CD14 (monocytes). (B) Different immune cells were examined for the presence of a signal for Exoria. Unstained MSC-EVs as well as a buffer control with Exoria were used as controls. (C) Analysis of the subcellular staining of following uptake of Exoria-stained EVs via imaging flow cytometry. Here, in addition to the light (bright field) and fluorescent microscopic images (PE channel; Exoria) merged images are shown.

## Discussion

Within this study we have evaluated the accuracy of dye mediated EV labelling as an example of well-studied MSC-EV preparations. Upon analyzing labelled MSC-EV preparations by IFCM we demonstrated that none of the conventionally used dyes, BODIPY-TR-CER, Calcein AM, CFSE and PKH67 allowed accurate labelling of MSC-EVs. In contrast, the novel dye Exoria allowed the quantitative labelling of EVs within our MSC-EV preparations without interfering with their immunomodulatory properties as monitored in a multi-donor mixed lymphocyte reaction assay. Furthermore, upon removing unbound dye by ultrafiltration, EV up-taking cells were identified. Notably, CD81^+^ EVs were stained more intensively with Exoria than CD63^+^ EVs, indicating potential differences in membrane compositions of both EV subtypes which significantly influences their stainability. In this context it is worth mentioning that upon comparing the intensity of Exoria labelled HEK293T and THP-1 EVs in an ongoing study, almost all HEK293T cell EVs could be efficiently labelled, while most THP-1 EVs remain unstained (data not shown). Thus, even though Exoria appeared as a very useful dye for EVs within our MSC-EV preparations, it should not be considered a pan-EV labelling dye.

For now, the EV-field is just started exploring the heterogeneity within given EV preparations with pioneering work performed by bead capturing approaches mainly against tetraspanins or other EV surface proteins (Koliha et al., 2016a; Koliha et al., 2016b; Kowal et al., 2016). Another critical component of EVs that affects dye incorporation is the lipid composition of EV subtypes. Many studies have demonstrated that EV preparations from different cell sources have varied lipid compositions (Skotland et al., 2019; Skotland et al., 2017). These differences have been studied as potential biomarkers for cancers such as colorectal cancer (Lydic et al., 2015), prostate cancer (Brzozowski et al., 2018) and Alzheimer disease (Su et al., 2020). Furthermore, many groups have utilized these differences to selectively capture EVs by utilizing different lipid binding molecules, i.e. chloral toxin B chain, Shiga toxin B subunit or Annexin V (Lai et al., 2016).

Whilst we were unable to identify the cause of the difference in EV subtype labelling, our study clearly demonstrates that experimenters need to critically (re)evaluate the appropriability of EV labelling dyes for their purposes. Using conventional technologies such as differential centrifugation protocols for the EV preparation or particle quantification devices, the specificity of the EV labelling dyes for the EVs of interest cannot be investigated (Simonsen, 2019). To this end, novel analysis devices are required, allowing EV analysis of dye stained and antibody labelled EVs. In addition to IFCM, other devices have entered the field allowing the colocalization of at least two different fluorescent labels on a single EV-sized object, including a novel generation of flow cytometers for nanoparticles such as the NanoFCM device (Tian et al., 2020) or nanoFACS (Morales-Kastresana et al., 2020), plasmon resonance devices with fluorescence detection units such as the NanoView device (Srinivasan et al., 2019) or novel direct stochastic optical reconstruction (dSTORM) devices such as the Nanoimager (Helmink et al., 2020) or the Nano Particle Tracking device in fluorescence mode. Indeed, elaborate analysis of EV labelling results performed on a NTA platform in the fluorescence mode has revealed discrepancies in the labelling of EVs with PKH. These results imply PKH uptake might not be connected to EVs (Dehghani et al., 2020; Lai et al., 2015). Upon characterizing of CFSE stained EV preparations of immature dendritic cells, CFSE was qualified as an appropriate labelling dye for these EVs (Morales-Kastresana et al., 2017b). Furthermore, EVs of different cell types differ in their esterase content, which effects the utility of CFSE labelling of generated EVs.

Overall, the results obtained in this and in other studies question the reliability of broadly used “EV labelling” dyes, challenging the interpretation of many EV studies which use dye-labelled EV preparations for the identification of potential EV target cells. For now, it is a common strategy in the field, to label EV preparations with “EV dyes” and perform uptake experiments with the dye labelled EV preparations. In our opinion authors need to re-evaluate whether their EVs were indeed specifically labelled and whether the cells that took up labelled particles are indeed the target cells of the EVs. As such, EV uptake experiments remain extremely challenging. To the best of our understanding, the EVs within our MSC-EV preparations were accurately labelled, and we are confident that the Exoria labelled EVs were specifically taken up by the various immune cell types. However, it remains an open question, which of the labelled EV subtypes the cells took up most efficiently, be it CD9^+^, CD63^+^ or CD81^+^ EVs. We also consider that the different immune cell types may have preferences for different EV subtypes. Although our study fails to provide these answers, we hope that it helps to sensitize EV researchers around the globe for the challenges associated with the identification of EV target cells. Indeed, issues are further complicated by the fact that not necessarily all EV target cells take up EVs to process their signal. At least a proportion of EV mediated intercellular interactions might follow the *kiss and run* principle that EVs bind to receptor platforms on cells, activate these platforms and are shed off, similar to what has already been shown for synaptic vesicles (Chanaday et al., 2019; Wen et al., 2017).

## Acknowledgment

We thank the Westdeutsche Spender Zentrale (WSZE) for providing bone marrow samples of healthy donors. We are grateful to the healthy blood donors whose cells were used in the mdMLR assay. This study was supported by funds of the European Union (ERA-NET EuroTransbio 11: EVTrust [031B0332B] and the European Union’s Horizon 2020 research and innovation programme EVPRO under grant agreement No 814495 and AutoCRAT under grant agreement No 874671. The materials presented and views expressed here are the responsibility of the authors only. The EU Commission takes no responsibility for any use made of the information set out).

## Contributions

T.T., M.S., P.F.J. and B.G. conceived and planned the experiments; T.T. carried out the experiments with assistance provided by O.S. and A.A.-J.; T.T., M.S., P.F.J. and B.G. analyzed and interpreted experimental results; T.T. and B.G. wrote the manuscript. All authors provided critical feedback and approved the final version of the manuscript.

