## Supplement file for "Imaging flow cytometry challenges the usefulness of classically used EV labelling dyes and qualifies that of a novel dye, named Exoria™ for the labelling of MSC-EV preparations"

### Supplemental Figures and Tables

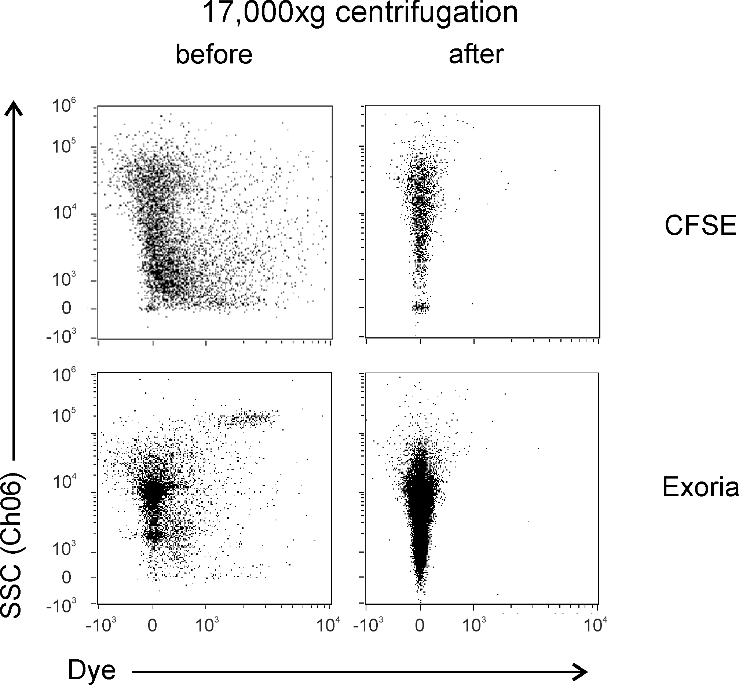

***Supplemental Figure 1: Centrifugation of the CFSE and Exoria dye reagent reduces background.*** *The staining solutions with CFSE or Exoria were spun at 17,000 x g for 10 min at RT to remove insoluble dye.*

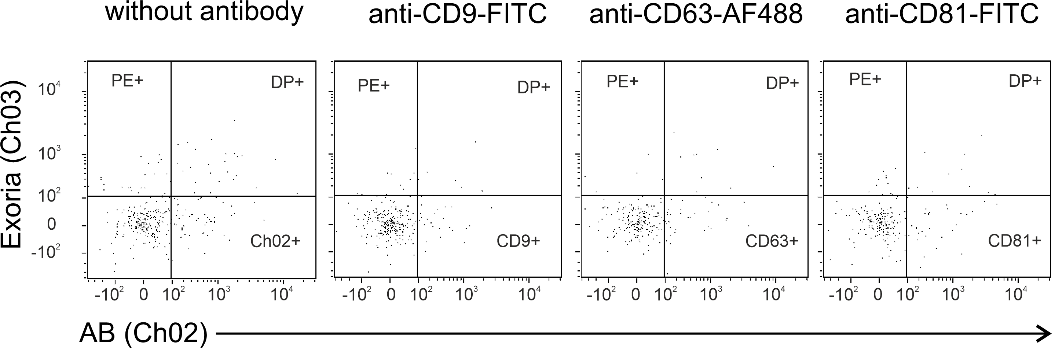

***Supplemental Figure 2: Exoria labelled EVs are detergent sensitive.*** *MSC-EV preparations stained with Exoria and/or anti-CD9, anti-CD63 or anti-CD81 antibodies following treatment with NP40. The non-NP40 treated reference plots are depicted in Figure 3.*

| **Antibody** | **Clone,**  **cat. #** | **Application** | **Manufacturer** | **Dilution** |
| --- | --- | --- | --- | --- |
| CD9-FITC (mouse) | MEM-61,  1F-208-T100 | IFCM | Exbio, Vestec, Czech Republic | 1:100 |
| CD9-PE (mouse) | MEM-61,  1P-208-T100 | IFCM | Exbio, Vestec, Czech Republic | 1:100 |
| CD63-AF488 (mouse) | MEM-259,  A4-343-T100 | IFCM | Exbio, Vestec, Czech Republic | 1:100 |
| CD63-PE (mouse) | MEM-259,  1A-343-T100 | IFCM | Exbio, Vestec, Czech Republic | 1:100 |
| CD81-FITC (mouse) | JS-64,  B25329 | IFCM | Beckman Coulter, Indianapolis, IN, USA | 1:100 |
| CD81-PE (mouse) | JS-64,  IM2579 | IFCM | Beckman Coulter, Indianapolis, IN, USA | 1:100 |
| Mouse IgG1-FITC | MOPC-21  555748 | IFCM | BD Biosciences, Heidelberg, Germany | 1:250 |
| Mouse IgG1-PE | MOPC-21  555749 | IFCM | BD Biosciences, Heidelberg, Germany | 1:250 |
| Mouse IgG1-AF488 | MOPC-21  557782 | IFCM | BD Biosciences, Heidelberg, Germany | 1:250 |
| Mouse IgG2a-FITC | S43.10  130-113-833 | IFCM | Miltenyi Biotec, Bergisch Gladbach, Germany | 1:250 |
| Mouse IgG2a-PE | S43.10  130-113-834 | IFCM | Miltenyi Biotec, Bergisch Gladbach, Germany | 1:250 |
| CD4-BV785 (mouse) | RPA-T4  300554 | FCM | BioLegend, San Diego, CA, USA | 1:50 |
| CD25-PE-Cy5.5 (mouse) | M-A251  555433 | FCM | BD Biosciences, Heidelberg, Germany | 1:50 |
| CD54-AF700 (mouse) | 1H4  A7-429-T100 | FCM | Exbio, Vestec, Czech Republic | 1:50 |
| CD8-BV650 | SK-1  344730 | FCM | BioLegend, San Diego, CA, USA | 1:50 |
| CD14-PO | MEM-15  PO-293-T100 | FCM | Exbio, Vestec, Czech Republic | 1:50 |
| CD19-ECD | J3-119  A07770 | FCM | Beckman Coulter, Indianapolis, IN, USA | 1:50 |
| CD56-APC (mouse) | HCD56  318310 | FCM | BioLegend, San Diego, CA, USA | 1:50 |

***Supplementary Table S1.*** *List of the antibodies and isotype controls used in this study, including their*

*clone and ordering number. The concentration of the isotype antibodies was adjusted to that of antibodies against EV-specific antigens.*

| Laser  [nm] | used Power  [mW] | max. Power  [mW] | Filter  [nm] |
| --- | --- | --- | --- |
| 375 | 70 | 70 | - |
| 488 | 100 | 100 | FITC (Ch02)  480-560 |
| 561 | 200 | 200 | PE (Ch03)  560-595  ECD (Ch04)  595-642 |
| 648 | 150 | 150 | APC (Ch11)  642-745 |
| 785 (SSC) | 70 | 70 | SSC (Ch06)  756-780 |

***Supplemental Table S2.*** *Applied laser settings for imaging flow cytometry.*

|  | Ch1 | Ch2 | Ch3 | Ch4 | Ch5 | Ch6 | Ch7 | Ch8 | Ch9 | Ch10 | Ch11 | Ch12 |
| --- | --- | --- | --- | --- | --- | --- | --- | --- | --- | --- | --- | --- |
| Ch1 | 1 | 0.029 | 0.042 | 0 | 0 | 0 | 0 | 0 | 0 | 0 | 0 | 0 |
| Ch2 | 0.051 | 1 | 0.05 | 0 | 0 | 0 | 0 | 0 | 0 | 0 | 0 | 0 |
| Ch3 | 0 | 0.13 | 1 | 0 | 0 | 0 | 0 | 0 | 0.02 | 0 | 0 | 0 |
| Ch4 | 0 | 0.064 | 0.49 | 1 | 0 | 0 | 0 | 0 | 0 | 0 | 0 | 0 |
| Ch5 | 0 | 0.017 | 0.155 | 0 | 1 | 0 | 0 | 0 | 0 | 0 | 0 | 0 |
| Ch6 | 0.015 | 0.02 | 0.04 | 0 | 0 | 1 | 0 | 0 | 0 | 0 | 0 | 0 |
| Ch7 | 0.023 | 0.003 | 0.003 | 0 | 0 | 0 | 1 | 0 | 0.015 | 0 | 0 | 0 |
| Ch8 | 0 | 0.032 | 0.008 | 0 | 0 | 0 | 0 | 1 | 0.012 | 0 | 0 | 0 |
| Ch9 | 0 | 0.004 | 0.084 | 0 | 0 | 0 | 0 | 0 | 1 | 0 | 0 | 0 |
| Ch10 | 0 | 0.002 | 0.041 | 0 | 0 | 0 | 0 | 0 | 0.084 | 1 | 0 | 0 |
| Ch11 | 0 | 0.001 | 0.012 | 0 | 0 | 0 | 0 | 0 | 0.025 | 0 | 1 | 0 |
| Ch12 | 0 | 0 | 0.003 | 0 | 0 | 0 | 0 | 0 | 0.013 | 0 | 0 | 1 |

***Supplemental Table S3.*** *Applied compensation matrix for imaging flow cytometry for PKH67 and Exoria labelled EVs and counter-stained with the antibodies.*
